# We might not notice a ‘mass’ extinction

**DOI:** 10.64898/2026.04.07.716927

**Authors:** Giovanni Strona, Corey J. A. Bradshaw

## Abstract

**Background:** There is overwhelming evidence that global change is having widespread, detrimental impacts on biodiversity. Population declines and local disappearances have been recorded with increasing frequency across all taxa, resulting in a steady rise in the number of threatened species. However, the number of documented extinctions remains counterintuitively low (∼ 1000 species across all kingdoms) compared to the sense of emergency pervading the scientific community. In isolation, that figure might fuel scepticism about the biodiversity crisis, but when put into context, it reveals that current extinction rates might be comparable to those that occurred during past mass extinction events estimated from the fossil record (≥ 75% extinctions within < 2 million years). Although this is an important clue supporting the claim that we might now be witnessing a new (‘sixth’) mass extinction, it falls short of definitive proof. The claim bears such high importance that it requires exceptionally solid foundations. However, our main aim was not to ascertain whether current extinction rates qualify as a new mass extinction event in progress. Instead, we examined the intersection of potential future loss scenarios and species discovery rates to address the fundamental question of whether and when we will be able to confirm a mass extinction is under way.

**Advances:** Our extrapolations suggest that the timing for a mass extinction to materialise (2,604–34,808 years from now at 75% diversity loss) is consistent with past mass extinctions (e.g., 12,000–108,000 years estimated for the Permian-Triassic extinction to unfold) under modern extinction rates (loss of 0.004%–0.053% of global species richness per year). We identify the minimum necessary conditions in which we could confirm a mass extinction under the full range of assumptions related to total species diversity (ranging from < 1.8 million to 163.2 million animal species) and discovery rates (e.g., ∼ 13,110 new animal species described per year as of 2026, with the number growing by ∼77 species per year), and the associated timeframe required. We show that there are many realistic future scenarios where we would fail to detect a mass extinction in progress.

**Outlook:** Based on available evidence, the rate of global biodiversity loss might already be consistent with the standard definition of a mass extinction. But even if true, current extinction rate estimates (20–8343 times background rates) would not necessarily imply a mass extinction is currently unfolding, because this claim can only be verified *a posteriori*. Our projections instead indicate that there is a high risk of not recognising a mass extinction as it unfolds — 49% across all parametrisations we explored. Furthermore, the temporal scale required for a mass extinction to materialise is orders of magnitude longer than relevant policy and legislative horizons, a mismatch that might appear to absolve today’s society of responsibility. In reality, the opposite is true — underestimating the likelihood of already being on a trajectory toward a mass extinction could have catastrophic consequences for future generations and historical accountability. Future generations will be forced to confront a world they perceive as normal, unaware of how much better off humanity could have been.

## Introduction

Many have argued that we might be in the middle of the sixth mass extinction (*1*–*6*). However, recent studies have called into question the relevance of the ‘sixth mass extinction’ paradigm for contemporary biodiversity loss (*7*–*10*). Although not dismissing the hypothesis that an extinction event is underway, Wiens and Saban (*7*) raised several compelling arguments about the lack of comparability between today’s extinction crisis and mass extinctions inferred from the fossil record, including undetected extinctions, biased estimates of extant and past diversity, and mismatches of temporal scale and phylogenetic resolution.

Quantitative estimates derived from the available information describing recent, documented extinctions suggest that current extinction rates are substantially higher than the background rate — that is, the ‘normal’ extinction rate expected for a non-mass extinction measured from the fossil record (*6*). However, obtaining an accurate quantification of the magnitude of the ongoing extinction process poses many challenges (*11*–*13*). Qualitative arguments about the massive anthropogenic impact on the biosphere, and how these might cause a mass extinction, remain largely speculative. In fact, it is easier to identify the causes of extinction (e.g., habitat destruction, climate change, invasive species, etc.) based on specific examples than to measure or project their global effects on biodiversity (*2, 14*–*16*). There is now convincing evidence that most extinctions might be triggered by ecological cascades (e.g., loss of pollinators following plant loss, or parasite extinctions following host extinctions) (*1*–*3, 17*–*19*). But despite rapid theoretical developments in network ecology, we still have limited knowledge of the complexities of real ecological interactions and the networks emerging from them (*20*–*23*). Inferring co-extinctions from ‘primary’ extinctions therefore remains a challenge. For instance, even in the case of obligate interactions such as those between parasites and their hosts, errors can emerge either from overestimating parasite extinction risk when the host’s range is incompletely quantified, or underestimating it by excluding fundamental ecological processes such as density feedbacks and life-cycle complexities (*17*). In some cases, species labelled ‘extinct’ are accidentally rediscovered (*24*), but missing co-extinctions of dependent species in general due to a lack of data is likely much more common (*25, 26*).

Not only is there limited information on who interacts with whom, but there are also incomplete data describing who is who. Focusing on multicellular animals (i.e., excluding bacteria, fungi, plants, viruses, etc.), there are ∼ 2 million described species so far (*27*). This accounts for only a fraction of total diversity on the planet (both described and undescribed species), but there is no consensus on how small that fraction is. Estimates of global species diversity (including described and undescribed species) vary widely, from a few million to hundreds of millions (*28*–*30*). This potentially huge knowledge gap implies that most current and future extinctions involving undescribed species might happen unnoticed, a risk that could be amplified by higher extinction vulnerability of poorly known taxa (*13, 31, 32*).

If we are in the midst of a mass extinction, yet cannot document the event, the implications would be enormous for future humanity. It would be the ecological equivalent of the holocaust never making it into the historical record. In the short or long term, the absence of evidence for a potential (and plausible) ongoing mass extinction could easily be interpreted as evidence of absence, thereby supporting the continuation of harmful practices and hampering conservation, mitigation, and restoration efforts. The possibility of this occurring raises fundamental questions about whether we will be able to collect relevant evidence, under what circumstances, and when. Here, we examine the available data and model potential future scenarios based on those to address not only the question of whether we are currently in a mass extinction, but also whether we will ever have the appropriate data to be certain.

### Confirming a mass extinction based on data, not models

Undocumented extinctions might be lost forever from the record (rare fossilisation and possible future palaeontological discovery notwithstanding) (*33, 34*), and focusing on them might permit an evaluation of the chances that a potential mass extinction will fly under the radar. A thought experiment reveals the limits in assessing whether we are currently in a mass extinction. The experiment assumes documenting 75% diversity loss (*5, 7, 9*) (an admittedly arbitrary threshold) compared to current diversity as the necessary condition to confirm a mass extinction, and seeks to identify alternative future scenarios where this would be feasible, while also drawing a tentative timeline for the phenomenon to unfold.

If and when cumulative undocumented extinctions exceed 25%, it would then be impossible to reach the minimal condition to confirm a mass extinction. In alternative scenarios where ≥ 75% of species are described before going extinct, thereby satisfying the necessary condition, it would be *virtually* possible to document a mass extinction. We use the term “virtually” because the condition will be necessary, but not sufficient. In fact, the sufficient condition would require not only cataloguing 75% of (initial) global diversity, but also documenting the extinction of those species. There is obviously a massive difference between the two processes in terms of effort and timeframe — although describing a novel species is a one-time task, documenting an extinction requires extensive and continued monitoring to collect satisfactory evidence of the species’ true disappearance. But at the very least, the discovery of a species means that it is on record, making its future absence potentially noticeable.

To do the thought experiment and assess the frequency of undocumented extinctions in the future, we need to combine (*i*) realistic estimates of global diversity, (*ii*) projections of global loss of diversity following current (and future) extinction rates, and (*iii*) future trends in species descriptions (estimated by combining recent description trends with alternative future scenarios accounting for technological development and cultural shifts). Given the uncertainty associated with all three aspects, we explored a range of scenarios covering most of the uncertainty ranges.

### Uncertainties in extinction rates and the temporal dimension of mass extinctions

Before determining if the biosphere is in the midst of a mass extinction, we first need to define this term. Any definition is arbitrary, but for simplicity, we can refer to the most commonly used definition: the loss of ≥ 75% of global diversity within a ‘short’ geological time (*5, 7, 9*) (< 2 million years, which represents < 0.4% of the time since the evolution of the Ediacaran biota (*35*)). So, a mass extinction is the outcome of a process that can last for a period that dwarfs the existence of *Homo sapiens*. It is therefore conceptually incorrect to use statements such as “we are in the middle of a mass extinction”, because the attribution works only *a posteriori*. Acceptable hypothetical statements are that we are experiencing extinction rates comparable to those that unfolded during previous mass extinction events, or that if current rates of diversity loss continue unabated, a mass extinction will eventually occur.

Numerically, the above threshold definition (75% loss over 2 million years) implies an average extinction rate of 0.00000069 species/global species richness/year (without considering speciation to replace lost species). If we consider both extinction and speciation under a simple birth–death model, the following exponential model can be used to derive the extinction rate:

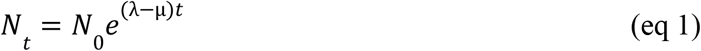

where *N*_*t*_ = number of species at time *t, N*_0_ = initial diversity, *λ* = speciation rate, and *μ* = extinction rate (both the latter expressed as species/global species richness/year). In the case of a hypothetical mass extinction where diversity decreases to 25% of *N*_0_ (*N*_*t*_/*N*_0_ = 0.25) after 2 million years, then:

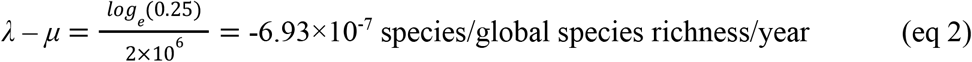

and

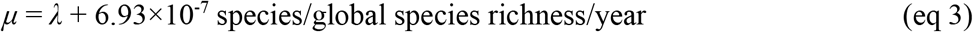

Assuming a representative animal speciation rate of *λ* = 10^-7^ year^-1^ (i.e., ∼ 1 speciation event lineage^-1^ 10 million years^-1^) (*36*), the corresponding extinction rate *μ* required to produce a 75% loss of diversity over 2 million years would be ∼ 7.93×10^-7^ year^-1^. This value corresponds to an annual loss of approximately 0.00000079 species/evaluated species/year (= 0.793 species million years^-1^), implying an average species lifetime of ∼ 1.26 million years under the assumption of constant rates. The estimated rate might be even lower if we consider a potential decline in diversification following global biodiversity loss and range contractions (*37*). How do these figures compare to the rates we can derive from inferred palaeontological or observed modern extinctions?

Different estimation methods give different estimates of extinction rates (values spanning ≥ 1 orders of magnitude). A widely used approach is deriving extinction rates from IUCN Red List data (*6, 38*) by comparing documented extinctions with monitored diversity, limited to species known to be extant in the modern scientific era (since ∼ 1500 Common Era). By including or excluding different categories from IUCN classification (e.g., with or without extant species labelled as in imminent danger of extinction), we can identify more or less conservative extinction rates (Fig. 1a,b). There are also published estimates derived from similar and different approaches, and for different taxa (Fig. 1c).

**Figure 1.**
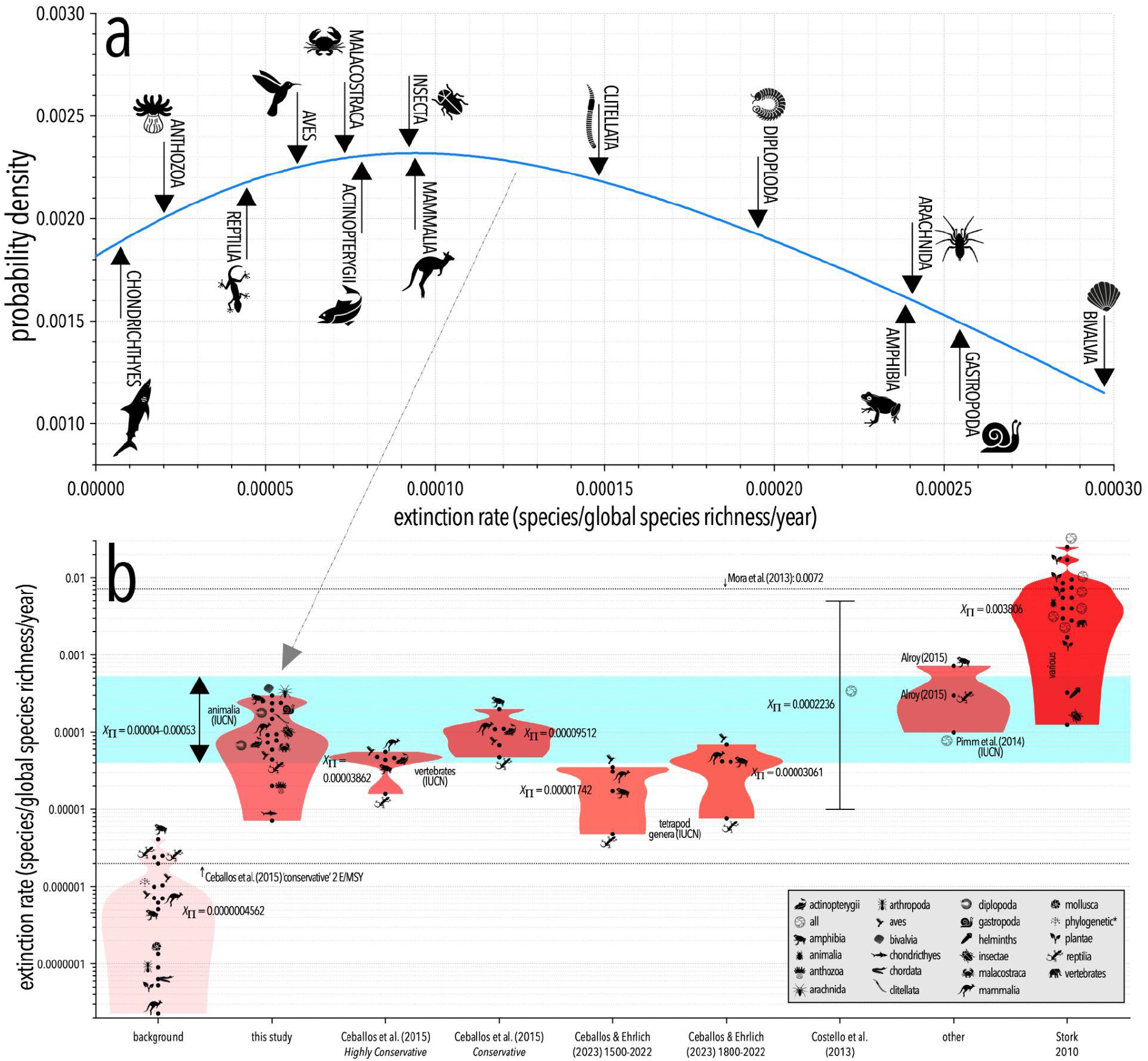
Extinction rate estimates (species/global species diversity/year) for different animal taxa and according to different sources (*6, 39*–*44*). (a) Probability density function across taxa for the ‘intermediate’ extinction rate derived from IUCN data for recorded extinctions (Extinct, Extinct in the Wild, Critically Endangered species also labelled as “possibly extinct” by the IUCN). Only classes with at least 100 evaluated species and a non-zero intermediate extinction rate are shown. (b) Combined information from a (range: 0.00004–0.00053 species/global species diversity/year calculated across all taxa; blue-shaded region) and the most cited and authoritative papers on the topic. We report both the background and current values per taxon by study, and violin plots show the density of aggregated extinction rates. Costello et al. (*40*) used values ranging from 0.01–5% decade^-1^ (i.e., 0.0001–0.005 year^-1^). The estimate used by Mora et al. (*41*) is 12–413 times higher than the IUCN-generated estimates.

If we take the geometric mean (X_Π_) of each grouping of extinction rates compared to the X_Π_ of the background estimates, current extinctions are 88–1162 times (this study), 38–209 times (Ceballos et al. (*6*) and Ceballos and Ehrlich (*6, 42*), respectively), 219 times (Pimm et al. (*43*)), and 8343 times (Stork (*39*)) higher than geometric mean background rates (Fig. 1c).

Compared to these values, the extinction rate estimated in the previous section giving a 75% loss of diversity over 2 million years (0.00000079 species/global species richness/year) is 51 and 674 times smaller than the lower and upper extinction rates estimated from the IUCN data, respectively (Fig. 1b,c) (ratio between extinct and evaluated species). In other words, current extinction rates are much higher than those required to precipitate a mass extinction. In fact, 0.00000079 species/global species richness/year is 2.5 times smaller than the estimated background extinction rate of 0.000002 species year^-1^ (i.e., 2 extinctions million species^-1^ year^-1^) used by Ceballos et al. (*6*). Thus, the 2-million-year window seems overly wide compared to available values. What would be a more realistic time frame?

To tackle the question, we can use the same model described in equation 1, and apply different estimates of extinction rate to derive a wide range of future projections of global biodiversity loss. The model assumes extinction and diversification rates will remain stable, while it ignores cascading ecological dependencies (*1, 3, 18*) and abrupt mass extinctions (*45*–*49*) that can accelerate species loss far beyond the predictions of a constant-rate exponential decay, hence likely providing an optimistic estimate of biodiversity loss.

As we outlined above and following Ceballos et al. (*6*), we can derive proxies for current (global) extinction rates by quantifying the ratio between the number of extinctions documented by the IUCN across all species for which a conservation status assessment is available over the last 125 years (i.e., since 1900). We estimated three alternative rates: (*i*) an optimistic scenario where we only included species classified by IUCN as Extinct or Extinct in the Wild; (*ii*) an intermediate scenario where we also included Critically Endangered species also labelled as “possibly extinct” by the IUCN; and (*iii*) a pessimistic scenario (i.e., with higher extinction rates) where we included all Critically Endangered species under the assumption that species at risk might contribute to an extinction debt (*50*–*52*) that could take its ultimate toll in the near future (i.e., decades). Resulting rates were 0.00004, 0.00011 and 0.00053, respectively (proportion of global species richness lost per year).

We then explored the timing for a mass extinction using 10 equally spaced extinction rates within this range, which resulted in an expectation of a mass extinction to be completed (diversity ≤ 25% of initial diversity) between the year 4,630 and 36,824 (median year = 9,917; mean year = 10,814; Fig. 2). In our simplified model, global extinction trajectories are only affected by extinction rates, not by initial diversity, because the probability of disappearing applies equally to all extant species (but clearly the raw number of extinctions would be proportional to global diversity). However, the prediction assumes current (2026) diversity starts at 100% global species richness, which is unrealistic because of extinctions that have occurred prior to today. Our estimates are therefore conservative.

**Figure 2.**
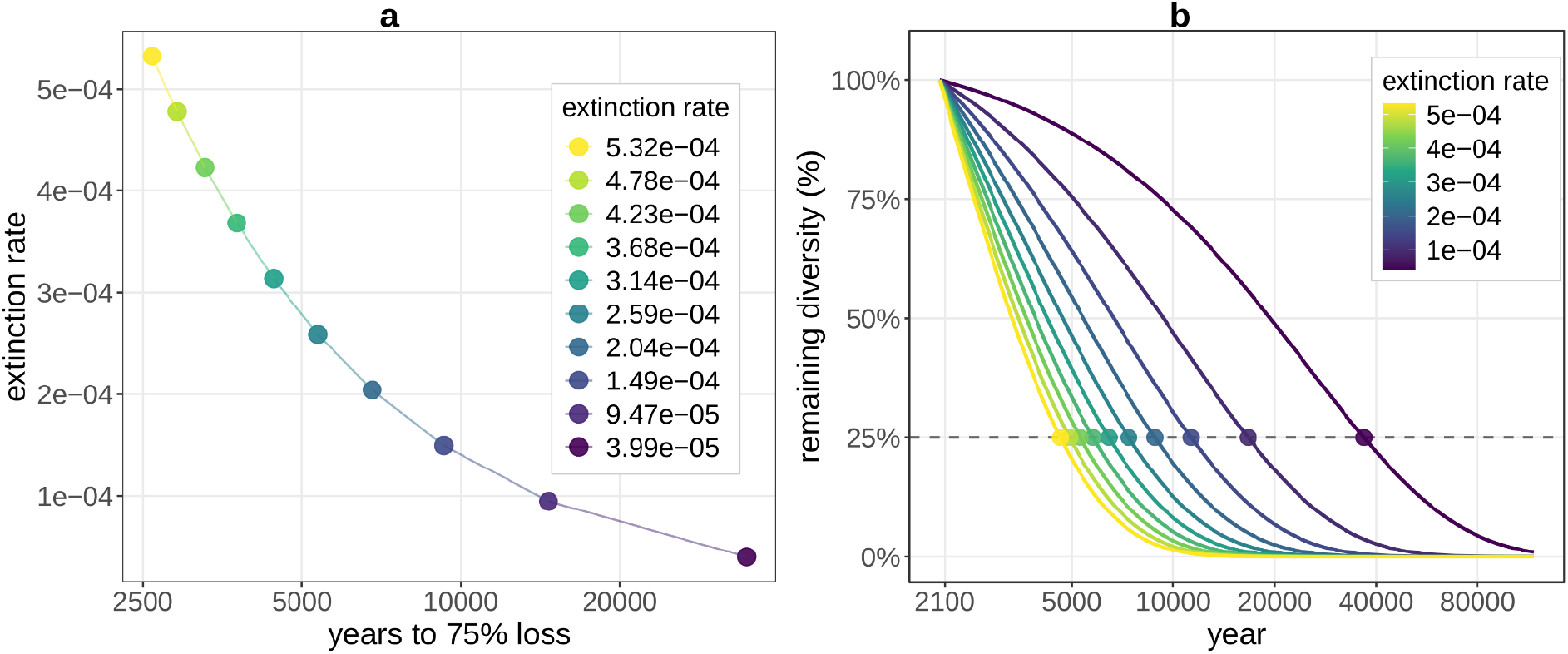
Time predicted for the disappearance of 75% of species modelled from equation 1 and using different extinction rates from ‘low’ and ‘high’ extinction rates estimated from IUCN data. Extinction rate is expressed as a proportion of total diversity lost year^-1^.

Comparing these numbers and past mass extinctions is complicated by the fact that the low temporal resolution of fossil data makes most mass extinction events appear instantaneous (*53*). A more precise estimate has been proposed for the Permian-Triassic extinction event 252 million years ago, with an estimated duration ranging from 12,000 to 108,000 years (*54*). Although these values are consistent in terms of order of magnitude with existing estimates of extinction rate, the corresponding lower and upper bounds are substantially lower than those we derived here, suggesting that the ongoing extinction process might be proceeding more rapidly than even the most catastrophic mass extinction event on record (*6*).

### Uncertainties in global diversity estimates

So far, around 2.2 million species have been described across the tree of life (*27, 55*). However, there is no consensus on how much extant diversity has yet to be described, especially for invertebrates and microorganisms. For the latter, even greater uncertainty is expected due to how challenges in detecting and cataloguing, as well as fundamental questions regarding species identity (*56*) (even for animals). Differences in estimates of global diversity from different sources span several orders of magnitude, from a few million to more than 100 million (Fig. 3). Such uncertainty does not affect estimates of relative global diversity loss and diversification (under the assumption that extinction and diversification rates apply equally to all species), but it has huge bearing on estimating description trajectories, because the number of species described per year depends primarily on study effort that is largely independent of total diversity (even though the likelihood of describing new species is expected to decrease with taxonomic knowledge — see below).

**Figure 3.**
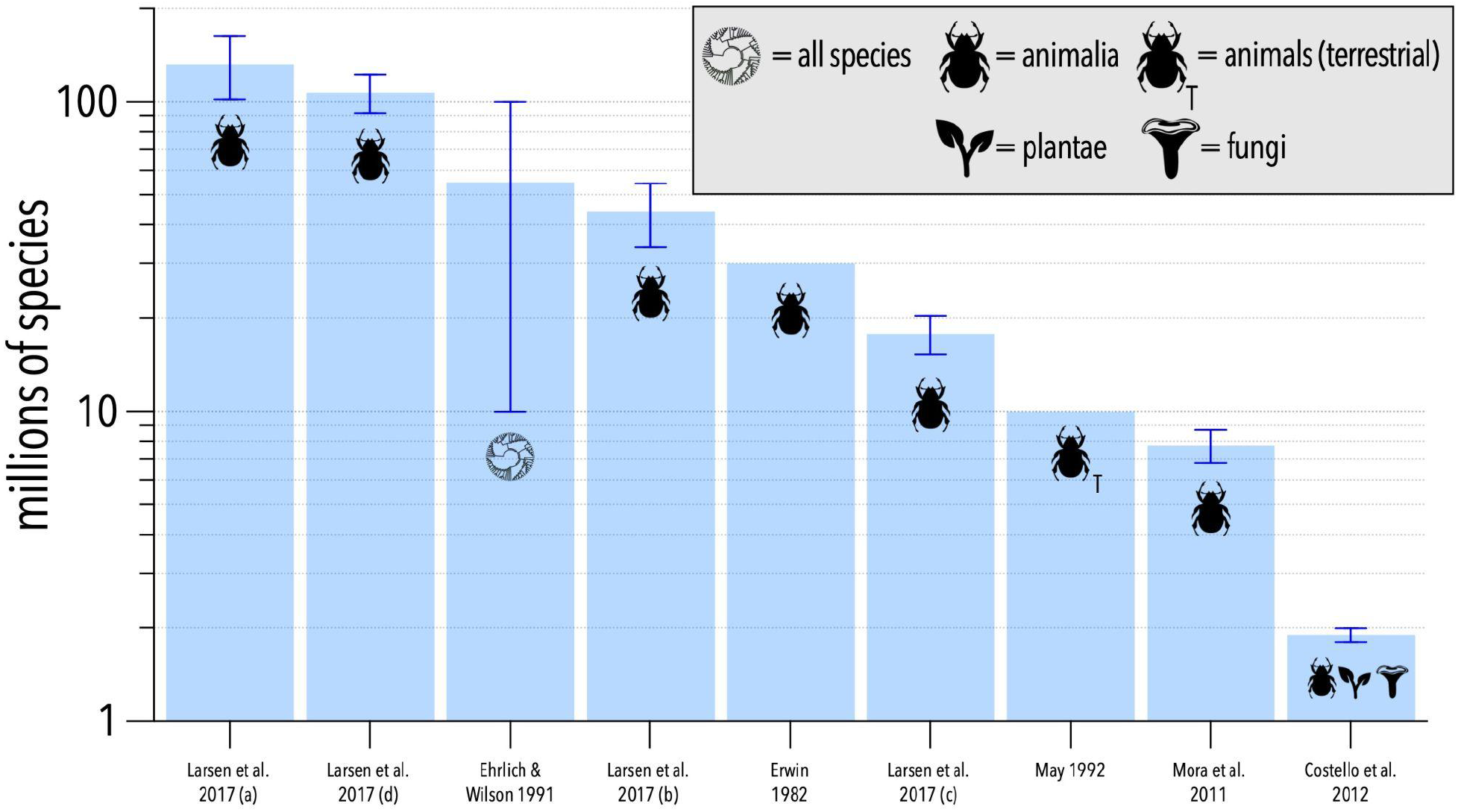
Various estimates of global species diversity (*28, 29, 57*–*61*). We sourced diversity estimates for the animalia where given (Erwin 1982; Mora et al. 2011; Larsen et al. 2017: 4 different estimates a–d corresponding to their Tables 1–4, respectively) (*28, 29, 57*) to compare to our estimates of animal extinction and discovery rates, but some estimates did not separate the animalia from other taxonomic groups (Ehrlich and Wilson 1991; Costello et al. 2012) (*58, 59*), or only provided a subset of the animalia (May 1988, 1992) (*60, 61*). Although there are many other estimates for groups at lower taxonomic resolutions (e.g., arthropods(*62*), mammals (*63*), etc.), we elected to show the range estimates from broader taxonomic groupings only.

### Uncertainties for future description rates

The rate at which species are (and will be) described depends on many phenomena, including interest in taxonomy, research effort, access to technologies, and technological development. In addition, what is considered a ‘described’ species in the future might not necessarily require peer-reviewed taxonomic descriptions, but only a deposited sequence (*64*), even though many taxonomists do not agree (*65, 66*). The rapid development of artificial intelligence might also create a system where classifications and even species descriptions might be done automatically (*67*), which could increase our ability to catalogue life (*68*). But interest in the subject might affect the rate of new species descriptions; for example, developments in the field might be hampered by a lack of dedicated effort and resources that are instead directed to other areas (*66, 69*–*71*). We might expect that the two processes could even balance the equation — an increase in description rates with improved technology might be offset by a decrease in interest, the emergence of other priorities, and/or consequent caps on allocated resources. Given the current situation where even applied science providing essential benefits to individuals and societies can be disrupted within days by reckless political decisions (*72, 73*), the long-term survival of basic sciences such as taxonomy appears increasingly endangered, likely relegated to a secondary role if not disappearing entirely (*74*).

Because of this complexity, estimating future description rates based on historical trends is uncertain, despite reliable data on the progress of taxonomy through time (from 1500 to now; Fig. S1). The recent trend in animal species descriptions (from 1970 to 2020) shows an approximately linear increase (with the number of descriptions per year growing by 77 species, and with > 13,000 species described in 2026), likely stemming from the benefits of novel technologies as well as increased access to education (e.g., a steady increase in PhD completions (*75*)), and the online revolution of the publication system (Fig. S2).

Considering these complexities and uncertainties, it is clearly impossible to be confident about the trajectory of future description trends. We therefore adopted a parsimonious assumption that current (positive) trends in species descriptions will continue steadily as long as a substantial proportion of extant diversity remains undescribed. We also assumed that discovering new species will become increasingly difficult as the pool of undescribed taxa diminishes (akin to a functional response) (*76*). Therefore, we propose a simple model that computes the number of future descriptions at year *y*_*i*_:

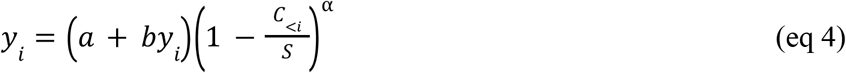

where *a* and *b* (8790.64 and 77.13, respectively) are the intercept and angular coefficients of a linear relationship between target year and number of species described in that year from 1970 to 2020, *C*_<*i*_ = total number of descriptions until *y*_*i*_, *S* = total estimated animal diversity (from the literature), and *α* is a parameter that modulates how the description trajectory slows as described diversity approaches estimated total diversity (*α* < 1 produces weaker slowing as undiscovered species become rarer; *α* > 1 produces more pronounced slowing). We explored different parameters for *α* (Fig. S3), but to avoid unnecessary complexity, here we report results only for *α* = 0.5.

### Thought experiment

We combined the global extinction model (equation 1) with future trajectories of species description to simulate the ‘race’ between taxonomic effort and biodiversity loss, and to estimate undocumented extinctions under different assumptions of current species diversity and extinction rate. We considered 100 unique combinations of initial global species diversity and extinction rate by taking 10 equally spaced values between the minimum and maximum diversity estimates (Fig. 1; i.e., 1.8 million–163.2 million), and 10 equally spaced extinction rates between our minimum and maximum IUCN-based estimate (0.00004–0.00053 species/global species richness/year). We ran each simulation from 2026 to year 10,000, modelling global diversity as a dynamic pool that is simultaneously depleted by extinctions and as species are catalogued by taxonomists. We assumed a self-limitation in the process of discovery, with undescribed biodiversity becoming increasingly rare as more species are catalogued (or go extinct) and are therefore more difficult to find.

The aggregated trajectories from all parametrisations (Fig. 4) show that on average, the loss of diversity exceeds the mass extinction threshold within the time frame of the simulations (10,000 years), ranging from < 25% of diversity loss to total annihilation. Undocumented extinctions ranged from ∼ 0% (for combinations of low extinction rate and low initial diversity; i.e., situations where the near totality of species had been already described) to > 50%.

**Figure 4.**
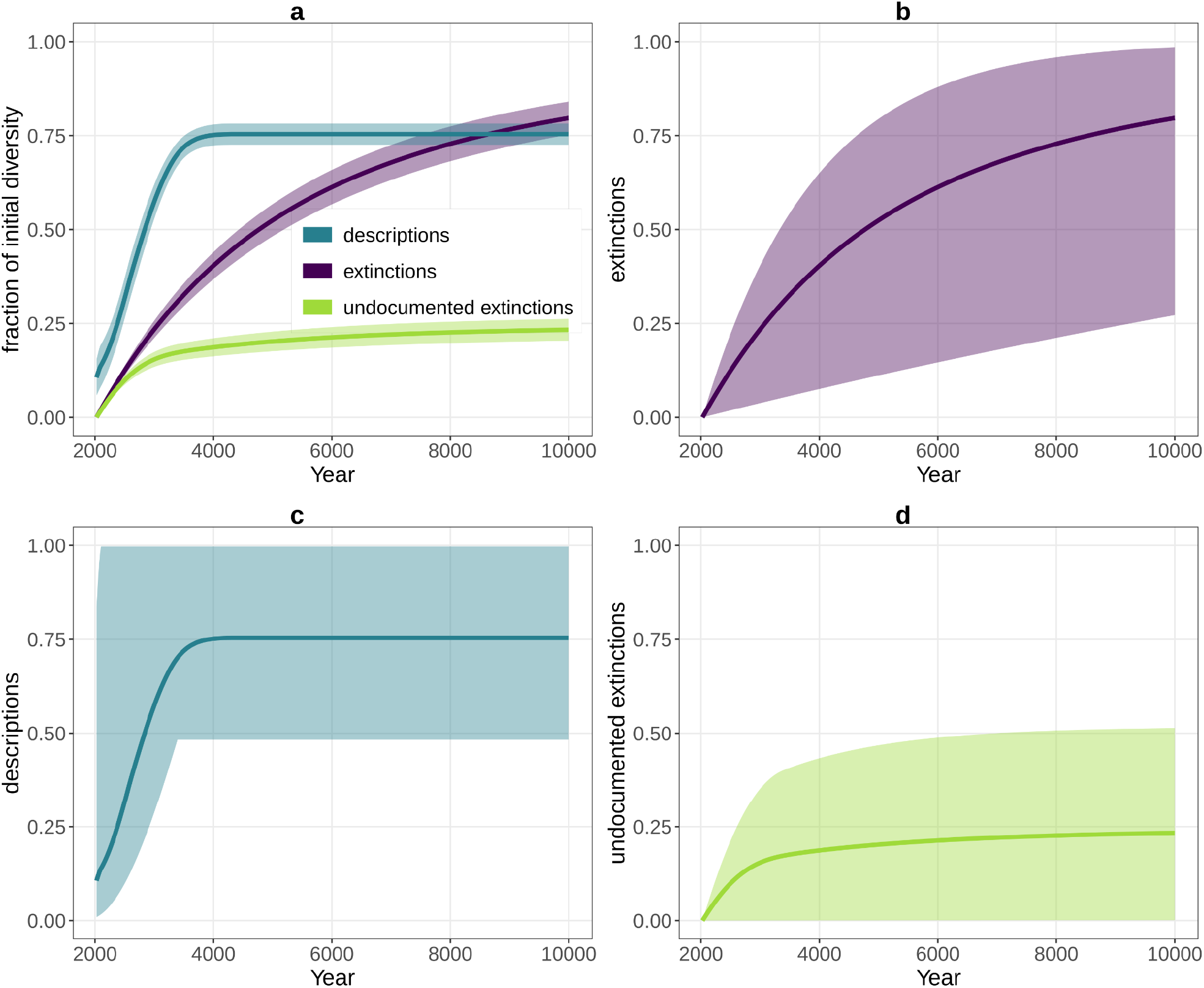
Description and extinction trajectories from the thought experiment under the full range of parameterisations. We tracked both total extinctions, descriptions and undocumented extinctions (i.e., species that went extinct without ever being described). We report values as a fraction of the initial diversity (which varied across simulations). Solid lines in all panels indicate average values across all simulations. Shaded areas indicate 95% confidence intervals in panel **a**, but they indicate the minimum-maximum range across all parameterisations in panels **b-d**.

In the thought experiment we assumed documenting at least 75% of global diversity (including extant and extinct species) was a necessary condition to confirm a mass extinction based on data. Because species descriptions will occur in parallel with extinctions, resulting in many species disappearing before discovery, the question is not only when we will be able to reach the 75% description threshold, but also whether — and under which circumstances — it will occur. Specifically, any scenario leading to ≥ 25% undocumented extinctions would make a formal, evidence-based conclusion of a mass extinction impossible (Fig. 5a,b). A fundamental determinant of whether we could discriminate between cases where proof is possible from cases where it is not is the global diversity estimate used. In scenarios with high (estimated) global diversity, describing most species would be increasingly challenging because diversity loss applies with equal probability to each individual species, but the number of species descriptions depends primarily on study effort and discovery capability/technology (even though diversity still affects description rates in the long run, in that it determines the slow-down due to saturation of described species; Fig. S3).

**Figure 5.**
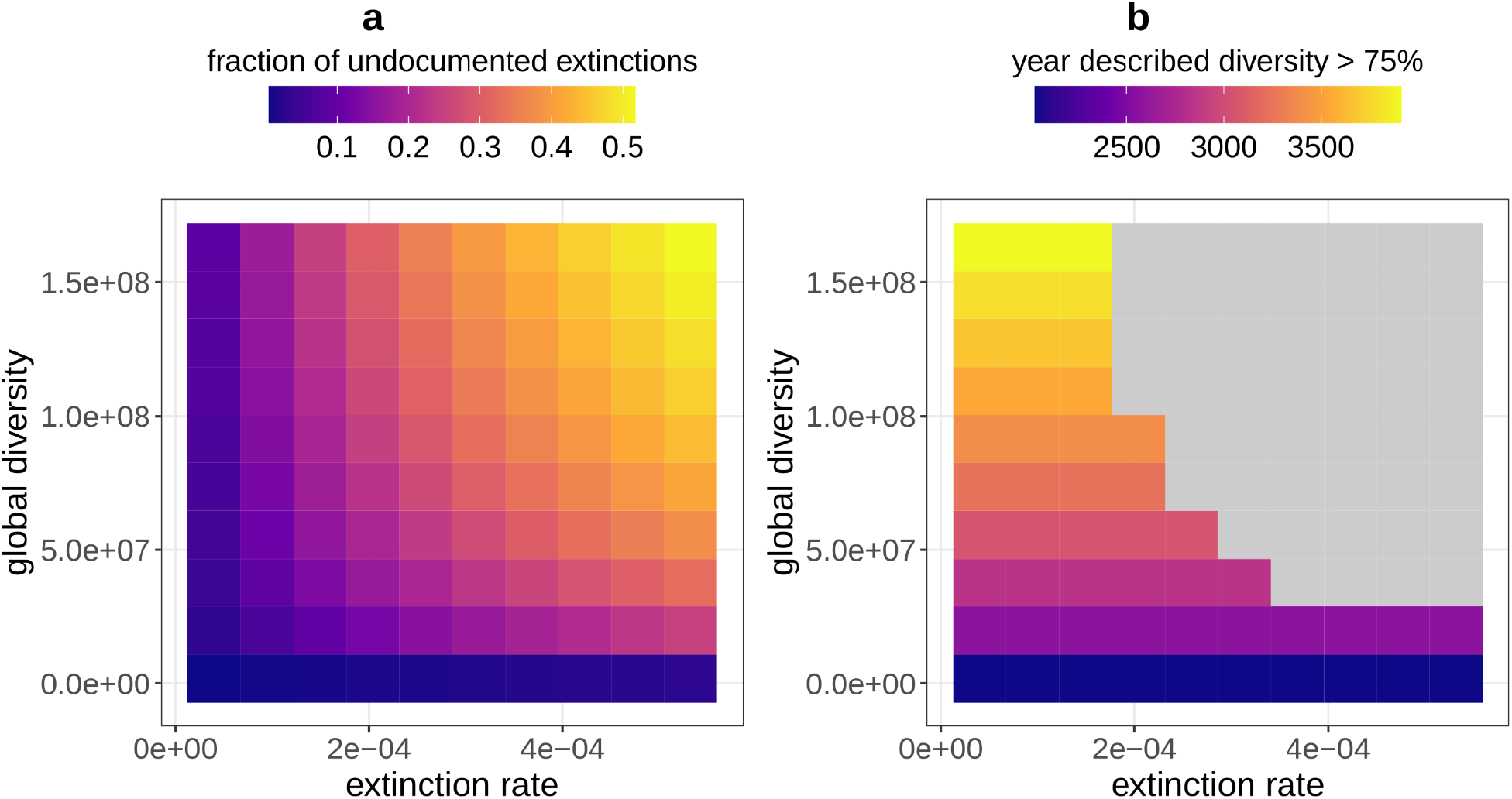
**(a)** Proportion of species that would go extinct unnoticed (compared to total, initial diversity) in our simulations. **(b**) Expected timing to describe 75% of current global diversity under different combinations of extinction rate and total global diversity estimate. The plot refers to a description rate curve with *α* = 0.5 (see Fig. 3). Grey cells identify all the parameter combinations that would result in not being able to achieve a 75% description (see Fig. 7).

In 49% of our scenarios (i.e., parameterisations), undescribed diversity exceeded 25%, making the documentation of a mass extinction impossible. Across these scenarios, the time undocumented extinctions surpassed the 25% threshold was, on average, in year 4231 (median year = 3143; lower and upper 95% confidence intervals = year 2616-year 10172). In the remaining scenarios where documenting a mass extinction was virtually possible, that possibility materialised, on average, in the year 2930 (median year = 2839; lower and upper 95% confidence intervals = year 2027-year 3911). Note that the latter scenarios, where it was possible to document at least 75% of global diversity before that is lost, are limited to the lower assumptions on global species diversity, i.e. they correspond to scenarios where most of Earth’s species have been already described. This also explains why the 75%-species-description target is expected, under those circumstances, to be met in a relatively short time (with an instant lower boundary). As soon as we move towards more realistic estimates of species diversity, the chances of a mass extinction to be documentable drop dramatically.

The time when ≥ 75% of species are lost would indicate the realisation of a mass extinction according to the most popular criteria. Undescribed extinctions will happen by necessity considering the initial knowledge gap, so the number of documented extinctions will be substantially smaller than the true number of extinctions. Avoiding models and extrapolations, and waiting until 75% of described species are observed to go extinct, we would document a mass extinction later than its true beginning.

### Underestimating the risk of unavoidable collapse

Most international biodiversity policies declare as an ultimate objective the halting of biodiversity loss (e.g., final text of the Kunming–Montreal Global Biodiversity Framework (*77*)). However, considering current ecological trends and their associated uncertainties, it is legitimate to question whether this objective remains achievable, and if so, how much time remains before meaningful intervention becomes impossible.

If we examine diversity trends from the fossil record, we can quantify how often episodes of diversity loss escalated to mass extinctions (in the lack of any anthropogenic influence). More generally, we can quantify how often a loss of any magnitude (e.g., 5 or 10% of global diversity) reached a target mass extinction threshold (e.g., ≥ 75%, or any other arbitrary threshold one wishes to apply). As a proof of concept, we analysed a time series of marine genera diversity spanning ∼ 500 million years to the present (*78*) (Supplementary Information). After removing long-term diversification trends using locally weighted regression, we identified episodes of sustained biodiversity decline relative to a dynamic baseline defined by recent diversity maxima. Each decline was treated as an independent crisis event, and its trajectory was followed until the point of maximum diversity loss. For every stage of each episode, we quantified the conditional probability that the observed loss would ultimately escalate to a mass extinction threshold (defined here as ≥ 75% diversity loss). We estimated these conditional escalation probabilities empirically using a monotonic generalised additive model fitted to the fossil data, allowing the probability of escalation to increase smoothly with the magnitude of diversity loss while remaining constrained between zero and one (see model details and statistics in Supplementary Information).

The resulting escalation model provides an empirical baseline probability that any biodiversity crisis ultimately escalates to mass extinction severity. For the fossil data we analysed, 6.27% of biodiversity loss episodes eventually reached a mass extinction threshold, with the probability of escalation growing rapidly once ∼ 30% of species have gone extinct (Fig. 6). Although this framework is only descriptive and cannot be used to make predictions, we can use it to contextualise our thought experiment and reveal the urgency of the current biodiversity crisis. In terms of escalation potential, the time frame for declaring a biodiversity ‘emergency’ is considerably shorter than the time required to achieve a mass extinction (Fig. 6). Even a baseline escalation probability (> 5%) would translate to an unacceptably high risk of eventually becoming a mass extinction when judged according to criteria used in engineering, aviation, or public safety (e.g., accepted risk of structural failures in buildings ranges from 0.01% to 0.001% year^-1^).

**Figure 6.**
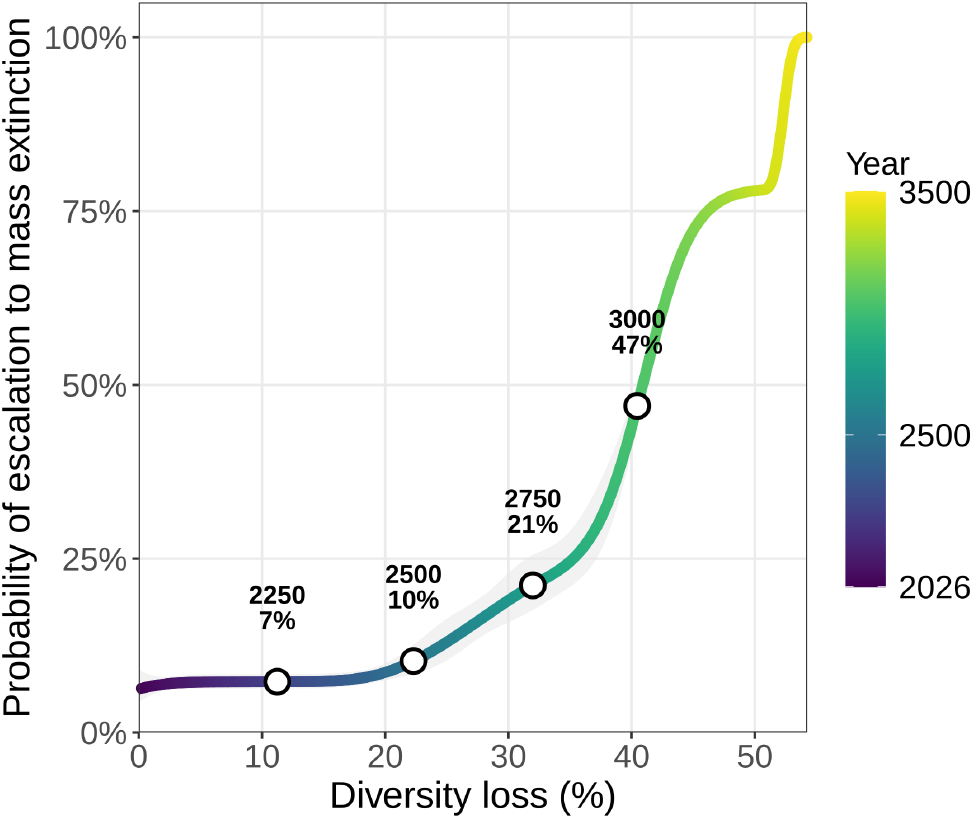
Probability of escalation to a mass extinction event as a function of biodiversity loss (%). Curves are based on a monotonic generalised additive model fitted to a time series of fossil marine genera diversity spanning ∼ 500 million years to the present (*78*), with shaded areas representing 95% confidence intervals. Coloured lines indicate the probability trajectory over time, with the colour gradient corresponding to calendar years starting from 2026. Markers highlight projected probabilities for selected future years under a high-extinction-rate scenario, derived from IUCN-based extinction rate estimates (see Supplementary Methods).

Most people have strong behavioural responses to or exhibit irrational fear towards highly improbable events. For instance, there are < 1 shark attacks/million people/year (*79*), and the probability of dying in a commercial flight is 0.00000017 (∼ 1 in 6 million flights (*80*)). Regardless, most people have an irrational fear of sharks (*81*), and many have a flying phobia (*82*). But no rational person would accept the individual risk of shark attack or dying in a plane crash if it were > 1 in 20. Current policies appear to accept large risks of biosphere collapse.

## Discussion

Although our analysis reveals the substantial uncertainty in claiming that we are currently in the midst of a sixth mass extinction, we have simultaneously demonstrated that current extinction rates are consistent with, and perhaps even worse than, those inferred for previous mass extinctions. However, the time required to achieve the commonly applied threshold of ≥ 75% diversity loss as a mass extinction according to our conservative model is in the order of thousands or tens of thousands of years from now. Depending on which assumptions of the magnitude of total global diversity, as well as any future patterns of extinction and description rates, there are many realistic scenarios (49% of our simulations) where we could pass a mass extinction threshold without ever noticing that we had.

We have also demonstrated that assumptions regarding the shape of the extinction loss curve have important implications for how a mass extinction might unfold. Under the simplified assumption of a stable extinction rate, the trajectory of global diversity loss is non-linear (exponential decay), meaning that extinction proceeds faster during the early phase of a mass extinction than towards its completion. This emphasises that preventing more extinctions earlier in the event compared to later is a more effective approach to limit the ultimate damage than being complacent and assuming future interventions are sufficient. This is especially important considering the potential breakdown of ecological networks precipitating even faster rates of co-extinction (*3, 18, 25, 26*).

Whether we can ultimately document a mass extinction is an interesting theoretical thought experiment, but the timeframe required for this to occur, even applying optimistic assumptions, renders any ensuing policy debates immaterial because of the obvious temporal mismatch between political choices made today and the fate of global biodiversity thousands of years from now. Although this mismatch could be used illegitimately to stifle a sense of urgency regarding the conservation of extant biodiversity, we instead argue that the undeniable evidence of extinction rates being consistent with mass extinctions, as well as the arbitrariness of threshold definitions, emphasise the severity of the current crisis. Even if the unfolding of a mass extinction might last thousands of years, the process leading to that endpoint is as much, or even more, important than the endpoint itself. Any progress of extinction exceeding the background rate should be considered undesirable from *inter alia* ecosystem (*83*), evolutionary (*84*), economic (*85*), and human health (*86*–*88*) perspectives. Therefore, even if ongoing biodiversity loss takes thousands if not tens of thousands of years to qualify as a mass extinction, and even if we are not able to document it empirically, it would be a mistake to dismiss the problem as irrelevant now and pass it on to future generations simply because it does not align with short-term policy timeframe.

If we adopt an optimistic perspective, the ongoing biodiversity crisis and global extinction risk might be halted at some point. However, a common oversimplification inherent in existing biodiversity-related policies is that the biosphere is simply an aggregation of individual species. In reality, ‘biodiversity’ is an emergent result of how species are connected by complex networks of direct and indirect ecological interactions (*23*). All these complex linkages involve non-linearities and unpredictability in how the disappearance of any species might affect any other. In turn, this means that global diversity loss might be dominated by an accelerating feedback loop that could escalate to cause non-reversible collapse of a functioning biosphere. In addition, there is an important difference between *global* extinctions and *local* diversity loss (*3*) — the latter has obvious and immediate negative impacts, but is also part of the process leading to global extinctions(*89*). Under the assumption of ‘random’ loss and depending on spatial distribution, massive diversity loss is required to trigger complete species extinction (*90*). Therefore, even a small proportion of global extinctions might entail an enormous loss of local or ‘average’ diversity, as well as extensive range contractions, massive changes in community composition, and extensive loss of local ecosystem services.

In conclusion, there is a considerable risk that a mass extinction could unfold without being documented, with odds roughly akin to losing a coin toss. This oversimplification belies unmistakable collapse of local ecosystems. The relevant window of action therefore lies between the period of failing to notice the process and having it proceed until it is too late to limit the damage. Some could be lulled into the false sense that the temporal scale makes the problem irrelevant to daily life, despite the vast body of evidence linking the integrity of natural systems to the wellbeing of humans (*83, 85*–*88*). Like the frog failing to notice the gradual rise in temperature of the heated water in which it rests, the narrative should instead emphasise that failure to change our course now removes the option for survival later.

## Acknowledgements

C.J.A.B. thanks the European Commission Joint Research Centre (Italy) for financial support. C.J.A.B. acknowledge the sovereign Traditional Owners and custodians (First Nations) of the unceded lands, seas, and skies where he lives and works, including Kaurna in Tarndanya/Adelaide, Peramangk in Bukatila/Mount Lofty Ranges, and Ngarrindjeri Ruwe in Murrundi/lower Murray River, Kurangk/Coorong, and eastern Fleurieu Peninsula.

## Statistical software and reproducibility

We did all analyses and simulations in R (v. 4.5.3). Code available at github.com/giovannistrona/mass_extinction/ (doi: 10.5281/zenodo.19371503). A compressed archive including both the code and all data, ensuring full replicability of our simulations and analyses is available at https://doi.org/10.5281/zenodo.19371534.

## Supplementary Information

### Extended Methods

#### 1. Global diversity estimates

We reviewed existing literature on global diversity estimates (*28, 29, 57*–*63*), and extracted figures for animal diversity. Those estimates ranged from 1.8 million to 163.2 million, with the largest number made up primarily by undescribed invertebrates. The smallest estimate is practically equivalent to the true number of known animal species (1,522,652 as of 2026 according to the Catalogue of Life (*27*); see next section). This means we effectively assumed that the most optimistic scenario regarding taxonomic completeness corresponds to a situation where practically all animal species have already been described.

#### 2. Descriptions

To explore trends in animal species description through time, we sourced all records from the Catalogue of Life “NameUsage” dataset (*27*). We retained only records with an accepted taxonomic status at the species rank, restricted to kingdom Animalia. We assigned year of first description using a priority-based rule applied to three Catalogue of Life date fields: (1) col:*basionymAuthorshipYear* (year species was described for the first time, under its original name); (2) col:*publishedInYear* (year of publication in which the species description appeared); and (3) col:*combinationAuthorshipYear* (year a taxonomist reclassified the species into a different genus, creating a new name combination). We used the first non-missing value in this sequence as the description year, prioritising the original description date where available. The combination year, used only as a last resort, reflects a taxonomic reclassification rather than a first discovery, and might therefore marginally overestimate the true date at which a species first entered the scientific record. We excluded records with no valid date across all three fields. We also excluded records with description years prior to 1500 CE or after 2025 (treated as data entry errors), as well as a large number (141,302) of extinct species (mostly from the fossil record). This yielded 1,504,924 valid animal species with a year of description.

From these data, we obtained annual animal species description counts from 1970 to 2020 (we excluded most recent years to remove biases due to the potential lag between actual descriptions and their appearance in Catalogue of Life), which we used to calibrate a linear model for the temporal trend in descriptions, obtaining the following equation (R^2^ = 0.86, p < 2×10^-16^):

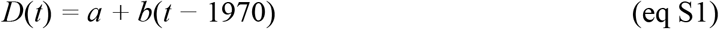

where *D*(*t*) is the number of species described in year *t, a =* 8790.6, and *b* = 77.1. The model therefore predicts 13,110 new species described per year (in 2026), with the number increasing by 77 species year^-1^ (Fig. S1).

Based on current knowledge, it is impossible to predict whether this trend will remain stable, increase (e.g., because of novel technologies facilitating species discovery and classification), or decrease (e.g., because of declining interest in taxonomy due to the rise of other priorities reducing resources allocated to basic sciences, a situation that zoology is already facing). Here we took the parsimonious assumption of no change in the future description rate, with the only caveat that as taxonomic knowledge approaches completeness, discovery rates must decline because fewer undescribed species remain. We modelled this saturation effect (functional response) by adding a depletion term to equation S1:

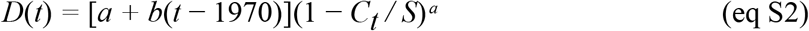

where: *D*(*t*) = number of new species described in year *i, a* and *b* = intercept and slope from the linear regression, *C*_*t*_ = cumulative number of described species up to year *t, S* = total estimated global animal diversity (described + undescribed), and *α* = depletion exponent governing the shape of the slowdown as undescribed species become rarer.

When *α* < 1, the slowdown as undiscovered species become rarer is less severe; when *α* > 1, it is more pronounced. We explored values of *α* from 0.0 to 1.0 in increments of 0.01 (Fig. 5a in main text). For all primary results we present in the manuscript, we used *α* = 0.5. In each simulation year, we capped new descriptions at the total number of remaining undescribed species to prevent the described pool from exceeding total estimated diversity.

#### 3. Extinctions

We obtained species-level IUCN Red List assessments for all animal species from iucnredlist.org (*55*). We excluded Data Deficient species from all analyses, retaining only species with a definitive threat category. For each species, we recorded the Red List category and whether it was flagged as Possibly Extinct. Following Ceballos et al. (*6*), we derived three estimates of extinction rate differing in their assumptions about which categories to count as extinct: (*i*) a conservative estimate counting only species classified as Extinct or Extinct in the Wild (EX/EW); (*ii*) an intermediate estimate additionally including Critically Endangered species flagged as Possibly Extinct (CR(PE)/CR(PEW)); and (*iii*) a more liberal estimate including all species listed as either EX, EW or CR . We focused only on recent human impacts, disregarding documented extinctions that happened before 1900. We expressed extinction rates (*E*) as the proportion of evaluated species lost per year (*S*_ev_), i.e.:

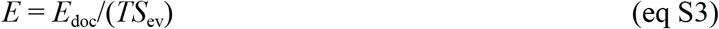

where *E*_doc_ = number of documented extinctions, *S*_ev_ = number of species evaluated, and *T* = time frame (125 years). The resulting rates were 0.00004, 0.00011, and 0.00053, respectively.

#### 4. Modelling undocumented extinctions

We combined description and extinction trajectories into a unified, annual-step simulation to track the ‘race’ between taxonomic effort and species loss. All simulations began in year 2025 and ran until year 10,000 (for trajectory plots) or until all thresholds were reached or up to year 100,000 (for threshold analysis).

We constructed a 10 × 10 factorial parameter grid, crossing 10 equally spaced extinction rates between our lower and upper IUCN-estimated rates (0.00004-0.00053 species/evaluated species/year), with 10 equally spaced total diversity values between the minimum and maximum of our diversity range (1.8 million to 163.2 million species). This yielded 100 unique parameter combinations.

For each parameter combination, the simulation tracked four state variables updated annually: div(*t*) = total number of extant species at year *t*, des(*t*) = cumulative number of species ever described at year *t* (whether or not still extant), des_alive(*t*) = number of described species still alive at year *t*, ue(*t*) = cumulative undocumented extinctions (species that went extinct before being described), and doc_ext(*t*) = cumulative documented extinctions (described species that subsequently went extinct).

We set initial conditions as follows: div(0) = total estimated diversity, varying across simulations (including described and undescribed animal species), des(0) = *C*_0_ (described species before 2025), des_alive(0) = *C*_0_ (all initially described species assumed to be alive in 2020), ue(0) = 0, and doc_ext(0) = 0.

For each subsequent year t, the updated equations were:

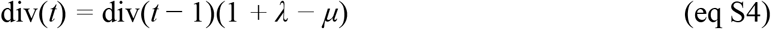

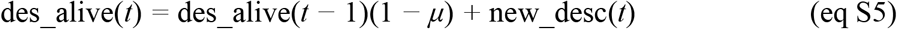

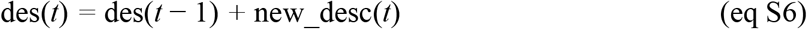

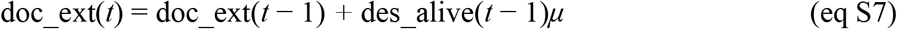

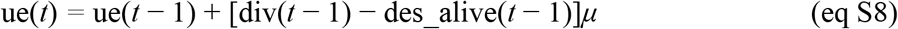

where new_desc(*t*) = the output of the depletion-corrected description model (eq S2), capped at the number of still-undescribed extant species: new_desc(*t*) ≤ div(*t* − 1) − des(*t* − 1). We fixed the speciation rate at *λ* = 10^-7^ year^-1^ throughout.

For each simulation, we recorded the first year at which each of the following thresholds was crossed: y_div25 = year when div(*t*) < 0.25*S* (i.e., 75% of species lost; mass extinction realised), y_des75 = year when des(*t*) > 0.75*S* (i.e., 75% of estimated total diversity described; necessary condition for evidential confirmation), y_ue25 = year when ue(*t*) > 0.25*S* (i.e., undocumented extinctions > 25%; necessary condition violated, confirmation impossible), and y_doc75 = year when doc_ext(*t*) > 0.75*S* (documented extinctions exceed 75% of total diversity). Simulations terminated early (before year 100,000) if they passed year 10,000 and all four thresholds had been recorded, to conserve computational resources. We truncated trajectories to year 10,000 for plotting. For each year, we computed across all 100 parameter combinations the mean, 95% confidence interval (mean ± 1.96 × standard error), and minimum, and maximum of each normalised state variable (proportion of *S*).

#### 5. Modelling risk of biodiversity loss escalation in the fossil record

We analysed genus-level marine invertebrate fossil data (*78*), first detrending diversity with a LOESS smoother (span = 0.75) and computing a rolling 25-bin maximum baseline to define relative diversity loss. We identified crisis episodes as contiguous intervals where diversity fell < 0.5% of the baseline; we recorded each episode’s nadir (maximum loss), and only used data up to the nadir for modeling. We added the start of each episode (0% loss) to anchor the probability of escalation. We fitted a monotonic binomial generalised additive model using the scam library in the R language, weighted by the number at risk, to estimate the probability of reaching a mass-extinction threshold (75% loss). The model provided an excellent fit to the observed escalation frequencies derived from the fossil record (edf = 5.96, χ^2^ = 70.08, deviance explained = 99.1%, adjusted R^2^ = 0.986, *n* = 77 observations). We then applied it to future projections of simulated global diversity trajectories under our three IUCN-based extinction rates to identify changes in risk of escalation to a mass extinction threshold (75% loss). In the main text we show the plot for the high extinction rate scenario, and here those for the intermediate and low rate scenarios (Fig. S4).

## Supplementary Figures

**Figure S1.**
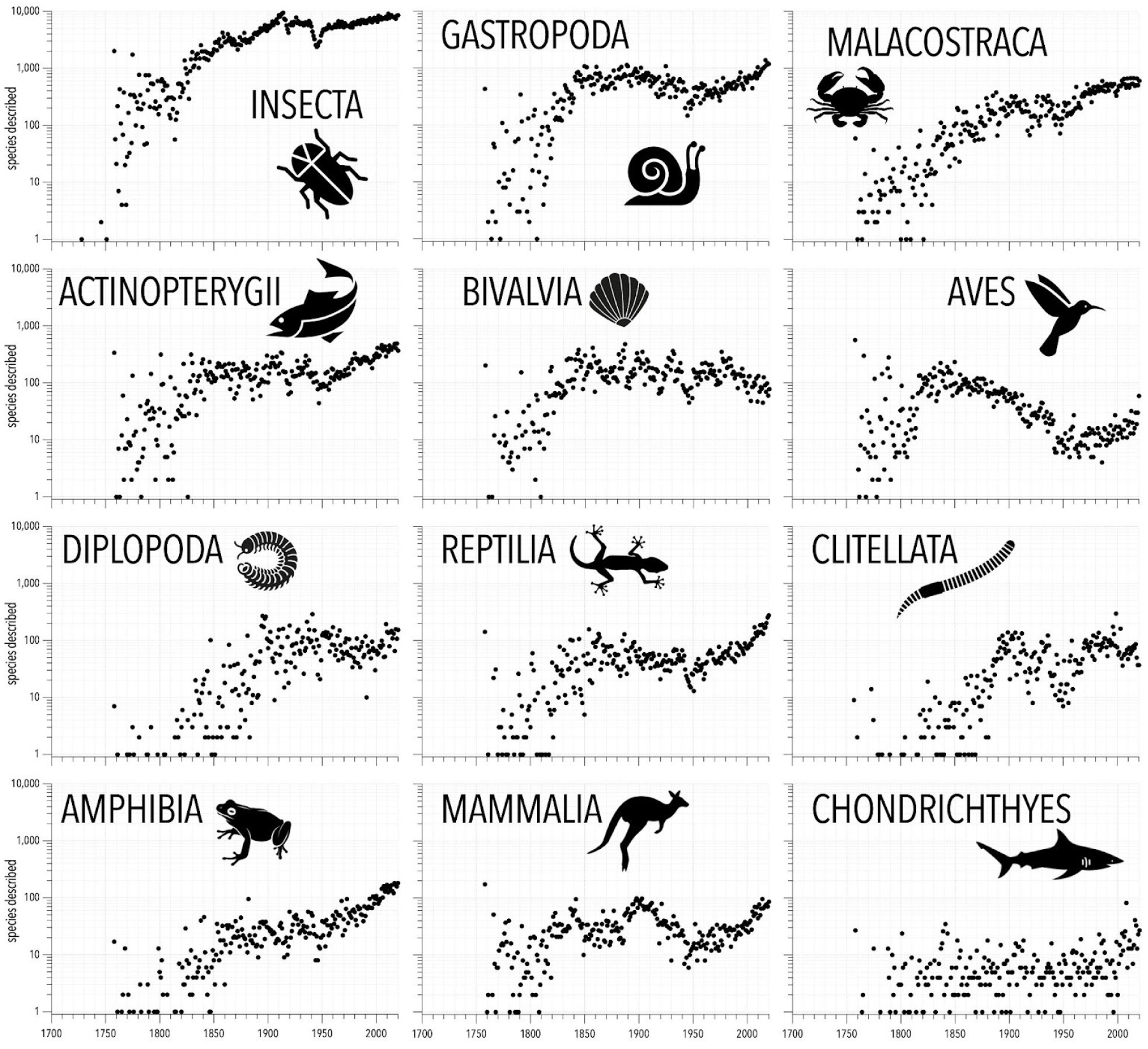
Temporal patterns in species description for different animal groups (log_10_ scale). Number of descriptions year^-1^ obtained by accessing year of first description from the Catalogue of Life (yearly extended version 2026) (*27*).

**Figure S2.**
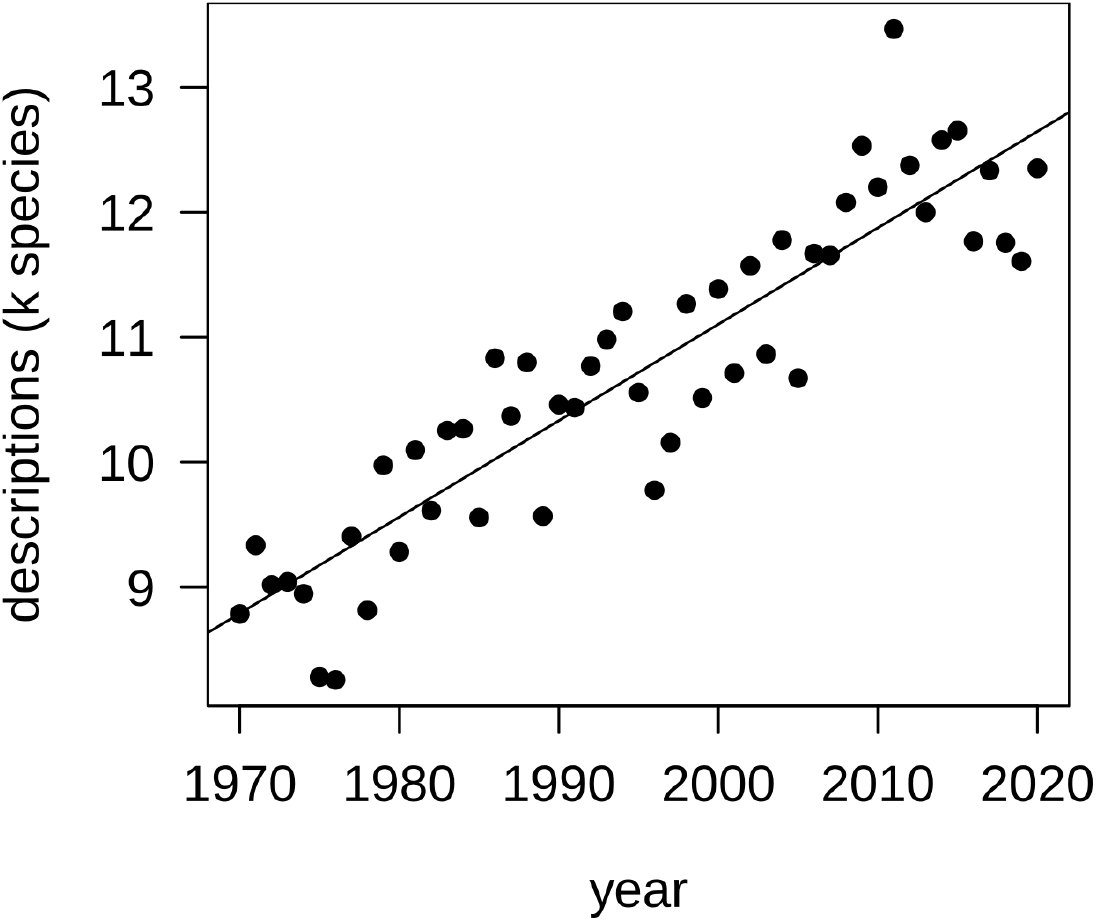
Trend in animal species descriptions in the period 1970–2020. Number of descriptions per year were obtained from the Catalogue of Life (*27*) (March 2026 yearly extended release). The solid line corresponds to the one described by equation S1.

**Figure S3.**
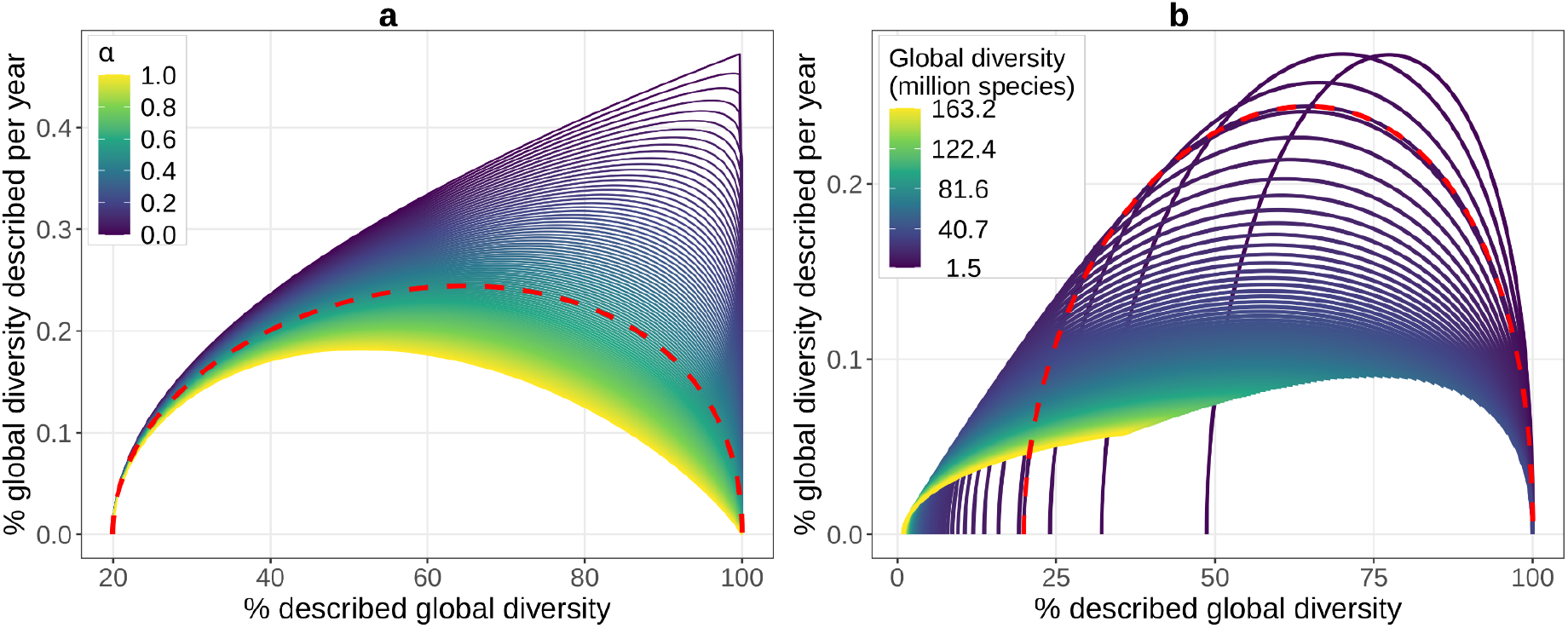
Hypothetical future trajectories of animal species descriptions according to different assumptions on how the rarity of undescribed species affects discovery rates (computed according to eq. 4). Both plots show how the fraction of total diversity described per year changes as taxonomic knowledge approaches completeness, in a hypothetical, extinction-free scenario. The starting number of species described per year (for 2025) is 13,033 species, and the starting annual linear increment (without considering *α*) is 77.1 species. Initial described diversity is 1,504,597 species. Panel (**a**) shows how description trajectories vary for different values of *α*, assuming a global total (described and undescribed) animal diversity of 7.7 million species(*29*). Panel (**b**) shows how trajectories vary under different assumptions of global animal diversity (*28, 29, 57*–*61*) for *α* = 0.5, that is the value we used in our thought experiment.

**Figure S4.**
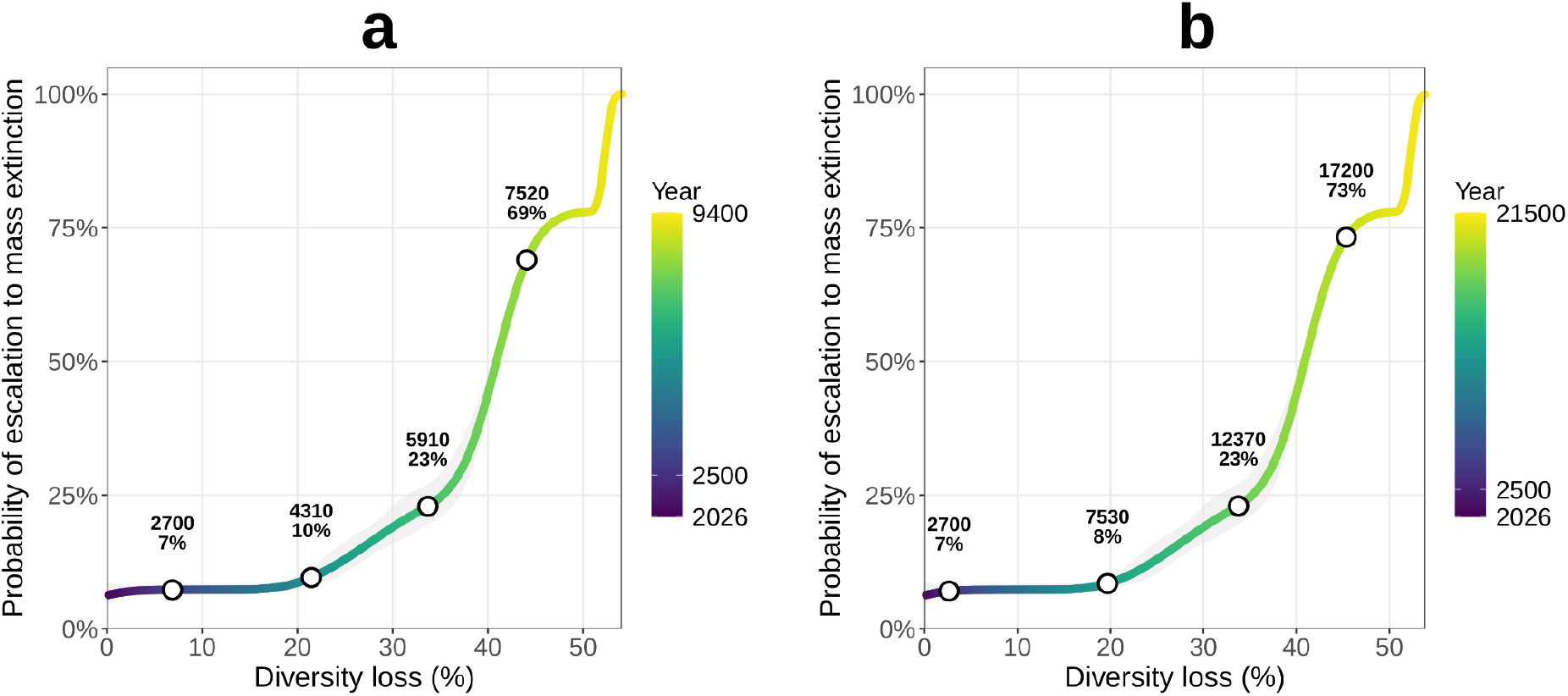
Probability of escalation to a mass extinction event as a function of biodiversity loss (%). The curves are based on a monotone generalised additive model fitted to a time series of fossil marine genera diversity spanning ∼ 500 million years to the present(*78*), with shaded areas representing 95% confidence intervals. Coloured lines indicate the probability trajectory over time, with the colour gradient corresponding to calendar years starting from 2026. Markers highlight projected probabilities for selected future years under an intermediate (a) and low (b) extinction-rate scenarios derived from IUCN-based extinction rate estimates (namely, 0.00011 and 0.00004).

